# Association indices for quantifying social relationships: how to deal with missing observations of individuals or groups

**DOI:** 10.1101/117044

**Authors:** William Hoppitt, Damien Farine

**Author notes:** Addresses for correspondence: William Hoppitt Damien Farine.

## Abstract

Social network analysis has provided important insight into many population processes in wild animals. Constructing social networks requires quantifying the relationship between each pair of individuals in the population. Researchers often use association indices to convert observations into a measure of propensity for individuals to be seen together. At its simplest, this measure is just the probability of observing both individuals together given that one has been seen (the simple ratio index). However, this probability becomes more challenging to calculate if the detection rate for individuals is imperfect. We first evaluate the performance of existing association indices at estimating true association rates under scenarios where (i) only a proportion of all groups are observed (group location errors), (ii) not all individuals are observed despite being present (individual location errors), and (iii) a combination of the two. Commonly-used methods aimed at dealing with incomplete observations perform poorly because they are based on arbitrary observation probabilities. We then derive complete indices that can be calibrated for the different types of observation probabilities to generate accurate estimates of association rates. These are provided in an R package that readily interfaces with existing routines. We conclude that using calibration data is an important step when constructing animal social networks, and that in their absence, researchers should use a simple estimator and explicitly consider the impact of this on their findings.

## INTRODUCTION

A foundation of animal social network analysis is estimating the frequency that two individuals associate or interact. Social networks are typically a description of interconnections that are formed by relationships (edges) among multiple individuals (nodes). Social network analysis is a set of tools that can be used to describe the patterns formed by these interconnections or evaluate these against hypotheses (Farine & Whitehead, 2015; Whitehead, 2008). One feature of social network analysis that is perhaps unique to studies on animal populations is that researchers rarely have a complete record of all interactions or all associations (but see Boogert, Farine, & Spencer, 2014; Farine, Spencer, & Boogert, 2015; Strandburg-Peshkin, Farine, Couzin, & Crofoot, 2015). Thus, relationships are often imperfectly sampled, which can introduce uncertainty in the social network. To account for variation in sampling effort and observation frequency, Cairns and Schwager (1987) outlined commonly-used association indices. These indices convert the number of observations of pairs of individuals seen associating or interaction into an association rate, representing their propensity to associate or their probability of being observed together.

Incomplete sampling of animal interactions or associations can occur due to a range of different reasons. We can classify datasets as having two possible types of missing data (Cairns & Schwager, 1987): (i) single or few observers can only collect data on one or a few groups at a time and miss many simultaneous associations or interactions occurring elsewhere, and (ii) individuals are difficult to observe or identify and missed even when they are present. In type (i), while a number of pairs of individuals (also known as dyads) are being observed together in one or more groups, the status of other individuals in the population is unobserved. In type (ii), when one or more groups are being observed they are incompletely sampled, resulting in data that suggests that certain dyads were not interacting or associating even when they were and could have been observed doing so. In both cases, the relationships inferred from the observed data is likely to be influenced by the amount of data that was missed. However, the propensity for each type of missing observations to impact our estimates of association or interaction rates and social network structure remains to be properly explored.

Properly controlling for missed observations is one of the most important steps in social network analysis. Using simulated data, Franks, Ruxton, and James (2010) identified the impact of missing observations when constructing social networks. They found that missing observations between known individuals was more problematic than missing individuals altogether, and concluded that social network sampling should maximize the amount of data collected about known individuals rather than maximizing the number of individuals sampled. One reason for this is because a key component of social networks, weak edges, are often disproportionately likely to be missed, and leaving these out can have profound implications on the structure of the social network (Granovetter, 1973). These findings are also supported by the work of M. J. Silk, Jackson, Croft, Colhoun, and Bearhop (2015) who explored the effect of completely missing individuals in the social network. They found that, with adequate sampling, having as few as 30% of individuals known can be enough to produce informative networks for hypothesis testing.

Missing observations that could have been recorded can have large impacts on the social network that is generated, and these impacts are made worse when particular individuals are missed more often than others. Farine and Whitehead (2015) recently demonstrated how small differences in the likelihood of observing individuals of different classes can introduce systematic biases in their social network. They first simulated observations of individuals associating with preferred and avoided associates. They then introduced a small observation bias, in this case reducing the probability of observing one of two classes of individuals to 80% by removing 20% of the observations of those individuals. This resulted in a significant effect of class on degree (the sum of the association strengths in the nodes with intact data was higher than in the nodes where data had been removed). This means that the social network estimated for the individuals in this population is incorrect.

In this paper, we theoretically re-evaluate existing association indices and derive new measures to deal with missing observations of groups, missing individuals in groups, and the combination of these. We show that the extent that existing association indices adjust estimates of association strength is entirely arbitrary, and are as likely to over-correct any bias that might occur as they are to reduce it. Existing association indices can also perform poorly at estimating relative association strengths, which has implications for many social network studies. We then derive improved association indices that enable researchers to correct properly for the biases arising from group location error and individual identification error, and discuss how to collect appropriate calibration data. Finally, we provide an R package “assocInd” that allows researchers to calculate accurate association indices for pairs of individuals from their observation data, and to simulate the effects of different types of errors on estimates of associations.

## THE SIMPLE RATIO AND THE HALF-WEIGHT INDEX

In many cases we wish to calculate an association index that estimates the proportion of time any two individuals, *a* and *b*, spend associated. Association indices typically range from 0 (the two individuals were never observed together) to 1 (the individuals are always seen together), and the association rates are used as a proxy to quantify the propensity for pairs of individuals to interact (Farine, 2015; Whitehead & Dufault, 1999), although the assumption that individuals interact in proportion to their association rate should be considered on a case-by-case basis (Castles et al., 2014). Association data is frequently collected by repeatedly sampling the population, and recording whom is observed in the same group in each sampling period. For any two individuals we can then calculate:

x the number of sampling periods with a and b observed associated
y_a_ the number of sampling periods with just a identified
y_b_ the number of sampling periods with just b identified
y_ab_ the number of sampling periods with a and b identified but not associated
y_Null_ the number of sampling periods with neither a nor b identified

In an ideal scenario, every individual is seen and correctly identified in every sampling period, such as in many captive populations, or at least we have the situation where y_Null_ = 0. Intuitively, in the ideal scenario researchers can validly use the simple ratio index (SRI), *x/*(*y_a_* + *y_b_* + *y_ab_* + *x*), as an estimate of the proportion of time A and B spend together. However, when errors arise from missing observations of individuals or groups, it is less clear that the simple ratio is appropriate. The most commonly-used approach for correcting association indices to account for missing observations is to reduce the weighting given to observations of just one individual (because we have a lower confidence in these). Because missing observations are widespread in behavioural research, many researchers use the half-weight index (HWI): 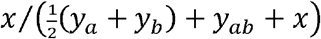 This index is believed to correct for the biases arising from such error, in particular when individuals are relatively more likely to be detected when they are apart than when they are together. When investigating the performance of association indices, Cairns and Schwager (1987) found that the HWI resulted in lower bias and lower error for a given estimate than the simple ratio when observations were missed. However, whilst this has served as useful justification for many researchers, it is also important to note that Cairns and Schwager (1987) reported up to 4 times greater error in the HWI than what they achieved using a maximum likelihood function (see also below). Further, they noted a number limitations of association indices arising from hidden assumptions.

Here we revisit some of the assumptions of the half-weight index. Notably, we show that the extent to which the half weight index adjusts estimates of association is entirely arbitrary, and is as likely to “overcorrect” any bias that might occur as it is to reduce that bias. Note that an alternative variant to the HWI, the twice-weight index (TWI) *x*/(2 (*y_a_* + *y_b_*) + *y_ab_* + *x*), is a monotonic function of the HWI and thus we do not investigate it in this paper. Ginsberg and Young (1992) previously raised the issue that the HWI and TWI use arbitrary weightings, and predicted that association indices will continue to be widely used. Indeed, the HWI is still the most commonly-used index in animal social network studies.

To address the need to properly correct for biases arising from group location error and individual identification error, we derive improved association indices that can be calibrated independently for each study. We start by addressing the impact of group location error before moving on to the effect of individual identification error, and finally the combination of the two.

## CORRECTING FOR GROUP LOCATION ERROR

Here we start with the assumption that if you see a in a group but do not see *b* in that group, you know *a* is not with *b*, and vice versa. This assumption will be valid if there is no individual identification error, i.e. all the individuals in a group that is located by the researcher will always be identified. However, uncertainty remains for all of the sampling periods in which we did not see *a* or *b*, since we do not know if they were together during such periods.

Let us denote the event that a and b are together in a sampling period as *ab* and the event that they are not together as *!ab.* The aim is to estimate the association between *a* and *b*, *a_ab_* = *p*(*ab*). We start by developing a maximum likelihood estimator (MLE) for a_ab_. In any given sampling period, the probability we see only individual *a* (i.e. not *b*), is given by:

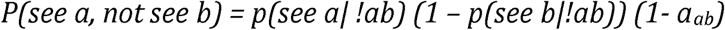

Note that *p(!ab) = 1-p(ab) = 1 – a_ab_.* The probability of seeing *a* and *b* in different groups is given by:

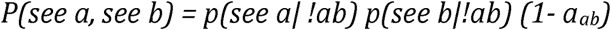

The probability of seeing a and b together in a group is:

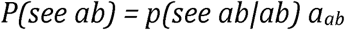

And the probability of seeing neither *a* nor *b* is:

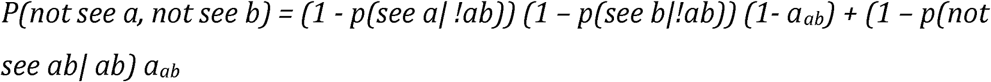

From this we can derive the log-likelihood, *L*, for the data obtained:

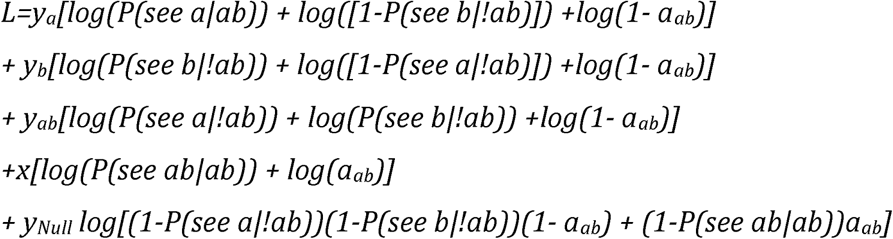

This simplifies to:

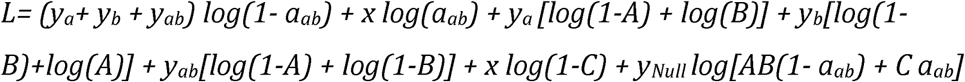

where *A = P*(*!see a/!ab*)*; B = P*(*!see b/!ab*)*; C = P*(*!see ab/ab*). We can find the maximum likelihood estimator, 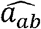 by solving:

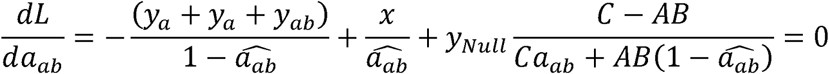

In practice *A* (the probability of not seeing a when a and b are not together), *B* (the probability of not seeing b when a and b are not together), and *C* (the probability of not seeing a or b when they are together) will not be known. Thus, to estimate an accurate value for the association strength between two individuals requires validation data at the level of individuals. However, progress can be made by making different assumptions about the relationship between *A, B* and *C.* First, if *C=AB*, the MLE is:

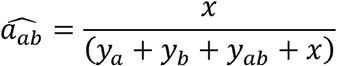

which is the simple ratio index. So, if the probability of failing to see *a* and *b* together is the same as the probability of failing to see both when they are apart, then the SRI is valid. Note also that the simple ratio is valid as an MLE if y_Null_ = 0, as intuition suggests.

Alternatively, we could assume that *C* = *AB*(1 + **ω**), where failing to observe a group containing *a* and *b* is more (*ω* > 0) or less (*ω* < 0) likely than failing to observe both the group containing *a* and the group containing *b* when *a* and *b* are not together. In this case the MLE is given by the solution to:

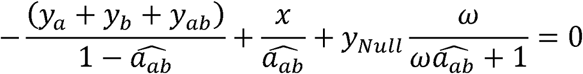

which can be re-arranged to form a quadratic:

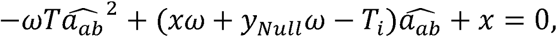

where *T_i_* is the number of directly informative sampling periods, i.e. *T_i_* = *y_a_* + *y_b_* + *y_ab_* + *x*, and T is the total number of sampling periods, i.e. *T* = *I* + *y_Null_⋅* The MLE is given by the lower root of this equation, i.e.

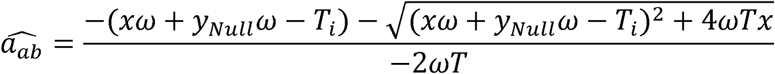

We term this index the group location error corrected index (GLECI). As expected, the GLECI reduces to the simple ratio index 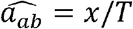 when *y_Null_* = 0. The standard error (see Appendix for derivation) can be calculated as for a proportion, 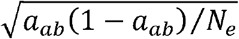 but with an effective sample size of:

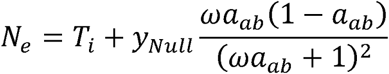

assuming sampling periods are sufficiently spaced in time and/or space to assume they are independent.

## CORRECTING FOR INDIVIDUAL IDENTIFICATION ERROR

Here we assume that there is no group location error, but define the probability of failing to identify an individual in a group that has been under observation, i.e. the individual observation error rate, as *ϵ*. We suspect this scenario will be rare, as individual identification error will usually be accompanied by group location error (see next section). However, we consider the case in order to analyse the effect of the two types of error. We get the following probabilities:

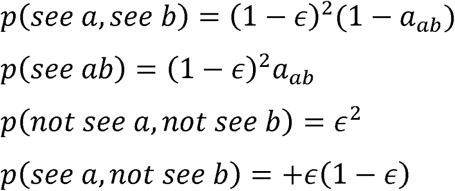

Therefore the likelihood for the data can be obtained as:

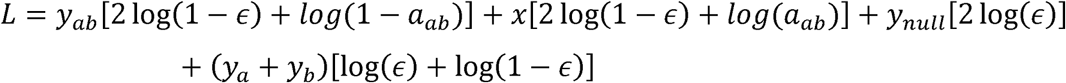

And the MLE found as follows:

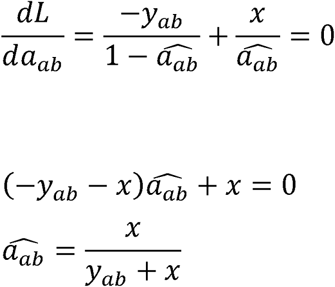

In this case the MLE 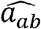 is a simple proportion, using only the *x* and *y_ab_* counts taken to the informative data, and requiring no calibration data. The terms *y_a_* and *y_b_* are not used, in contrast to the simple ratio and half weight indices, since they are known to be unreliable in this scenario: if only a is recorded, it is possible that b was associated with a and has been missed through individual identification error. We call this index the very simple ratio index (vSRI). The standard error (see Appendix for derivation) is calculated for a proportion as usual with 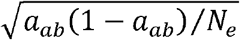 with an effective sample size of *N_e_* = *y_ab_* + *x*, assuming sampling periods are sufficiently spaced in time and/or space to assume they are independent.

## GENERAL ERROR MODEL

In practice, both types of error are likely to occur in a given sampling procedure. To model this situation, we define a model with a more general relationship between the errors in each count. We define:

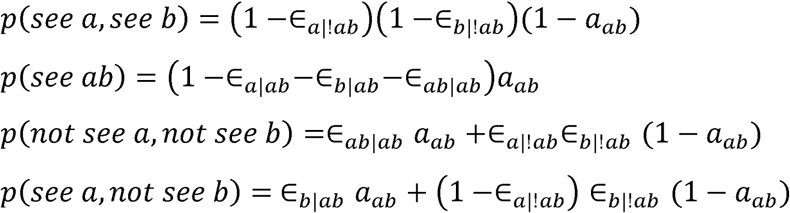

where ∈_*a|!ab*_ is the probability of missing *a*, given a is not with b; ∈_*a|ab*_ is the probability of missing *a*, but not b, given *a* is with b; and ∈_*a|ab*_ is the probability of missing a and b given they are together.

Note that the group location error scenario is the special case with ∈_*a|!ab*_ = *A;* ∈_*a|!ab*_ B; ∈_*ab|ab*_=*C*(1 + *ω*)*AB*=(1 + *ω*) ∈_*a|!ab*_∈_*b|!ab*_; ∈_*a|!ab*_=∈_*b|!ab*_=0. The individual identification error scenario is given by ∈_*a|!ab*_=∈_*b|!ab*_=∈ ∈_*a|!ab*_=∈_*b|!ab*_=∈(1–∈); ∈_*ab|ab*_=∈2.

Let us assume that the probability of missing both *a* and *b* when they are in the same group is (1 + *ω*) x the probability of missing *a* and *b* when they are not together (where *ω* > –1). The probability both *a* and *b* will be missed when they are together is therefore:

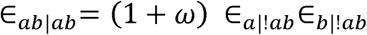

The probability of at least one of *a* or *b* being missed is ̀∈=1 – *p* (*see ab|ab*). Let us set ∈_*ab|ab*_ = ϕ ̀∈ where 0 < ϕ = 1. Since ̀∈ = ∈_*a|ab*_ + ∈_*b|ab*_ + ∈_*ab|ab.*_

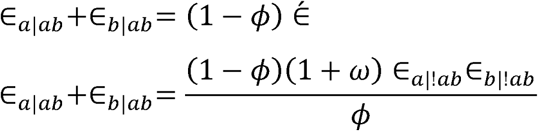

If we assume that ∈_a|ab_ /(∈_a|ab_ + ∈_b|ab_)= ∈_a|!ab_/(∈_a|!ab_ + ∈_b|!ab_) this gives us:

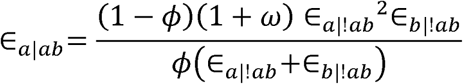

where *ϕ* determines the relative importance of group location error relative to individual identification error, with the group location error model given when *ϕ* = 1. We can now refine the probabilities given above:

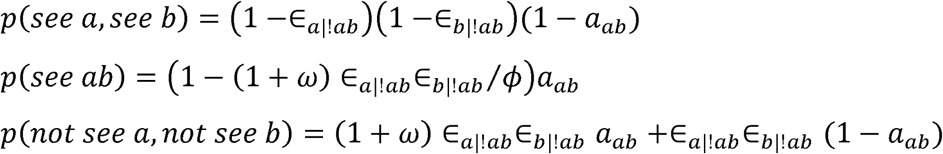

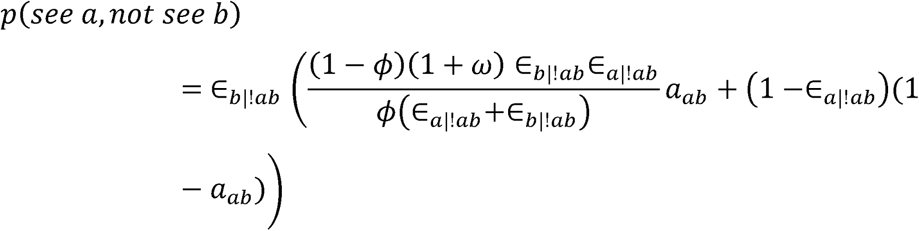

Giving a log likelihood of:

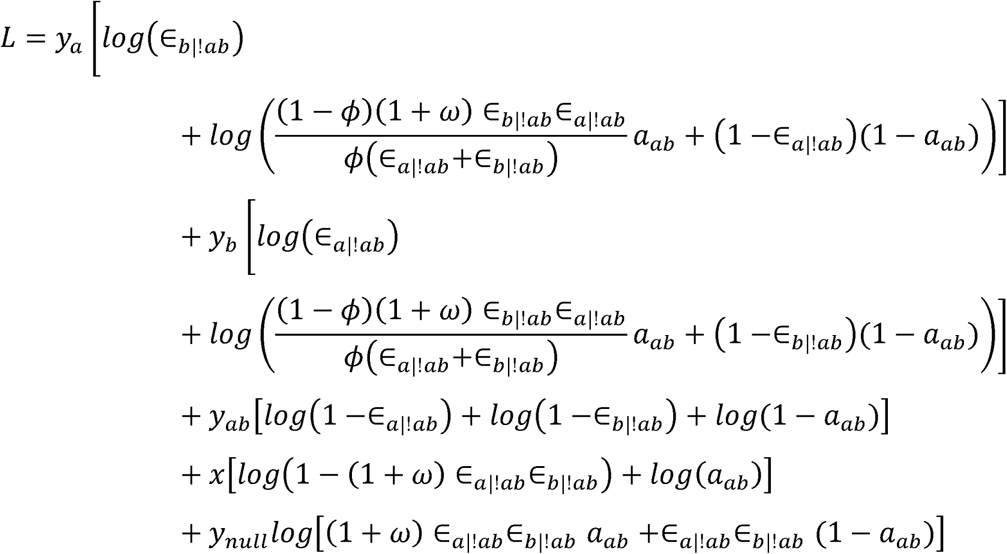

To obtain the MLE, 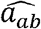 we need to solve the equation:

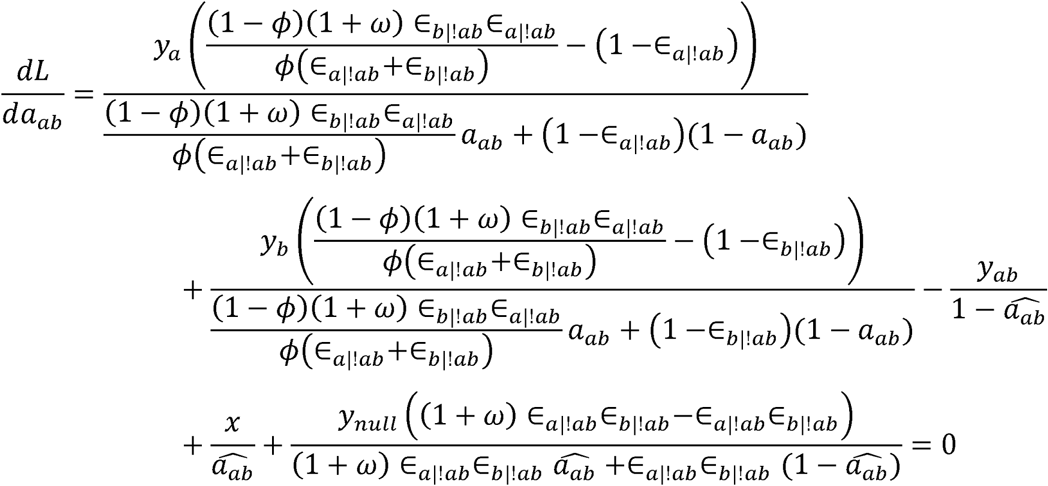

To generate an estimate of association using this function firstly requires values for *ω* (the group location error term) and *ϕ* (the error components importance term). These could reasonably be calibrated at the population level, i.e. we could assume that these quantities are constant across all dyads. However, the estimate also requires estimates for ∈_*a|!ab*_ and ∈_*a|!ab*_ which require calibration data at the level of individuals, making this approach infeasible in most cases. Nonetheless, we might obtain an approximate solution, 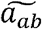 if we substitute a population averaged estimate (averaged across all dyads, or those dyads for which data is available) ∈= ∈_*a|!ab*_ = ∈_*b|!ab*_:

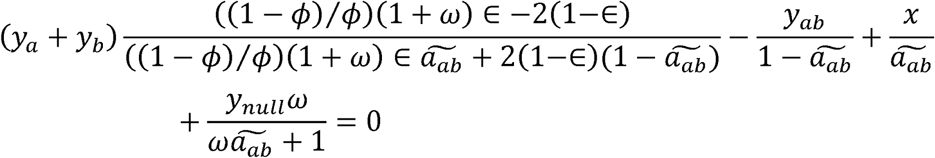

The calibration measures required to solve this equation are *ω*, *ϕ* and ∈, and can be solved using a non-linear equation solver. We call the solution to this equation, 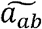 the combined errors index (CEI). In our R package, we provide a function that calculates the CEI in the R statistical environment (R Development Core Team, 2015), using the uniroot function in the rootSolve package (Soetaert & Herman, 2009). Note that by setting *ϕ* = 1, we reduce the model to the group location error model, giving 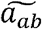 as the GLECI. We can also reduce the model to the individual identification error model by setting ∈_*a|ab*_ =∈_*b|ab*_ =∈(1–∈) and ∈_*ab*|*ab*_ ∈2, giving us *ϕ* = ∈/(2–∈) and *ω* = 0. Thus 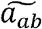 reduces to the vSRI.

## COMPARISON OF INDEX PERFORMANCE

In this section we examine how the SRI, HWI, GLECI, vSRI and CECI perform under scenarios where there is group location error, individual identification error, and a combination of the two. In simple cases we do this by first deriving expressions for the expected value of each index, and then dividing by the value it is intended to estimate, *E*[*I_ab_*]*/a_ab_*, thus showing us the circumstances under which *I_ab_* is biased upwards or downwards. However, we also recognise that in many circumstances only the relative sizes of *a_ab_* within the social network may be required, e.g. if estimating scale free node-based or network metrics. Consequently, we also derive (*E*[*I_ab_*]*/E*[*I_uv_*])/(*a_ab_/a_uv_*) to determine the circumstances under which each index tends to overestimate or underestimate ratios of association values. Here *u* and v denote a different dyad, so (*E*[*I_ab_*]*/E*[*I_uv_*])/(*a_ab_/a*_uv_) measures the bias when index *I_ab_* is used to estimate the relative strength of two associations.

In each case we also use simulations to illustrate the performance of the indices for a given set of values, investigate bias where we were unable to do so analytically, and examine the performance of Wald 95% confidence intervals calculated from the standard errors presented above. For each scenario, we simulated 10,000 datasets consisting of 1000 independent sampling periods for two individuals *a* and *b*, in the R statistical environment (R Development Core Team, 2015). We start by allocating the probability *a_ab_* = 0.5 that a and b were associating in a given sampling period. We then repeated all simulations with *a_ab_* = 0.25 and *a_ab_* = 0.75, and the results we qualitatively similar, so here we present the results for *a_ab_* = 0.5 only. We then simulated the observation process according to the models described above, to yield values for *y_a_, y_b_, y_ab_, x* and *y_nu_u* which we used to calculate the value of each of the target association indices. For each scenario, we repeated the simulation for a range of values of group location and individual identification errors. For the scenario including only group location error, we ran simulations for a range of values of *ω* = {–0.9, –0.8,…, 2.5} with A = B = 0.5, where A = P(!see a|!ab) and B = P(!see b|!ab). For the scenario including only individual identification error, we ran simulations for a range of values of ∈ = {0,0.05,…, 0.95}. For the scenario with both types of error, we varied *ω* = {–0.9, –0.8,…, 0.9}, ∈= {0.1,0.3,0.5} and *ϕ* = {0.1,0.3,0.5}, excluding impossible cases where(1 + *ω*) ∈*^2^/ ϕ* > 1, since this would mean there is a negative probability of observing a and b together. In each case we recorded the mean value for each association index, in order to detect bias, and the proportion of times the Wald 95% confidence intervals (calculated as ±1.96xSE) contain the true value for *a_ab_*.

However, recall that the CECI relies on an approximation, ∈= ∈_*a*|!*ab*_=∈_*b*|!*ab*,_ which replaces the individual specific error rates with population level ones. The simulations described above only test the performance of the CE when this approximation holds in the data, i.e. when error rates are the same across all individuals. Consequently, we re-ran simulations to test the performance of the CECI when there was individual variation in error rate. In each case we set the population mean error, ∈, arbitrarily to 0.5, but drew individual errors from a normal distribution with standard deviation *σ* = {0,0.2,… 2.0], discarding and resampling values that were < 0 or > 1, and likewise for ∈_*b*|!*ab*_ We then conducted the simulations as described above with *ϕ* = 0.5.

### Group location error only

We find that the GLECI is an unbiased estimator of *a_ab_* (see Table 1 and Fig. 1) across a range of group location errors *ω*. By contrast, the simple ratio is biased upwards when *ω* < 0 (i.e. when *a* and *b* are less likely to be missed when associated than both are to be missed when apart) and biased downwards when *ω* > 0. The commonly used HWI shifts the estimate of *a_ab_* upwards, such that it is biased upwards when *AB*(1 + *ω*) < (A + B)/2 and biased downwards when *AB*(1 + *ω*) > (*A* + *B*)/2. Consequently the HWI is only unbiased when the probability of seeing a and b together is equal to the average of the probability of seeing each of them apart. In our terminology, this is denoted (1 – *C*) *=* ((1 – *A*) + (1 – B))/2, giving *C* = (*A + B*)/2. Thus under this scenario, the HWI assumes that the probability of missing *a* and *b* when they are together is equal to half the probability of missing either *a* or *b* when they are apart. This seems to us to be an arbitrary a priori assumption, without the functionality to adjust the assumption using supporting calibration data.

**Figure 1:**
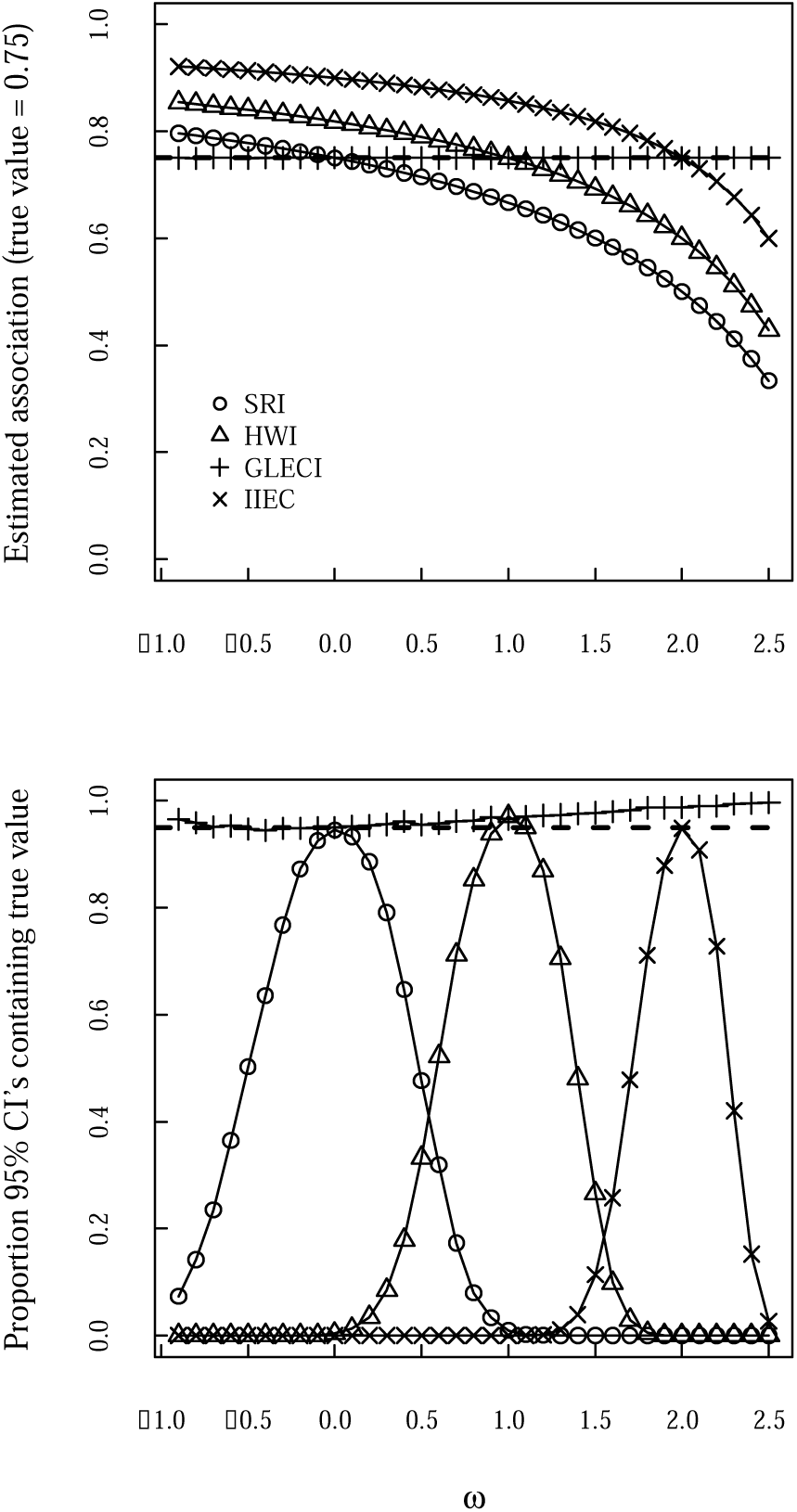
a) Bias in different association indices as a function of group location error (*ω*) when applied to simulated data; b) performance of 95% Wald confidence intervals as a function of *ω*. Similar results were obtained for a true association value of 0.25 and 0.75.

**Table 1:**
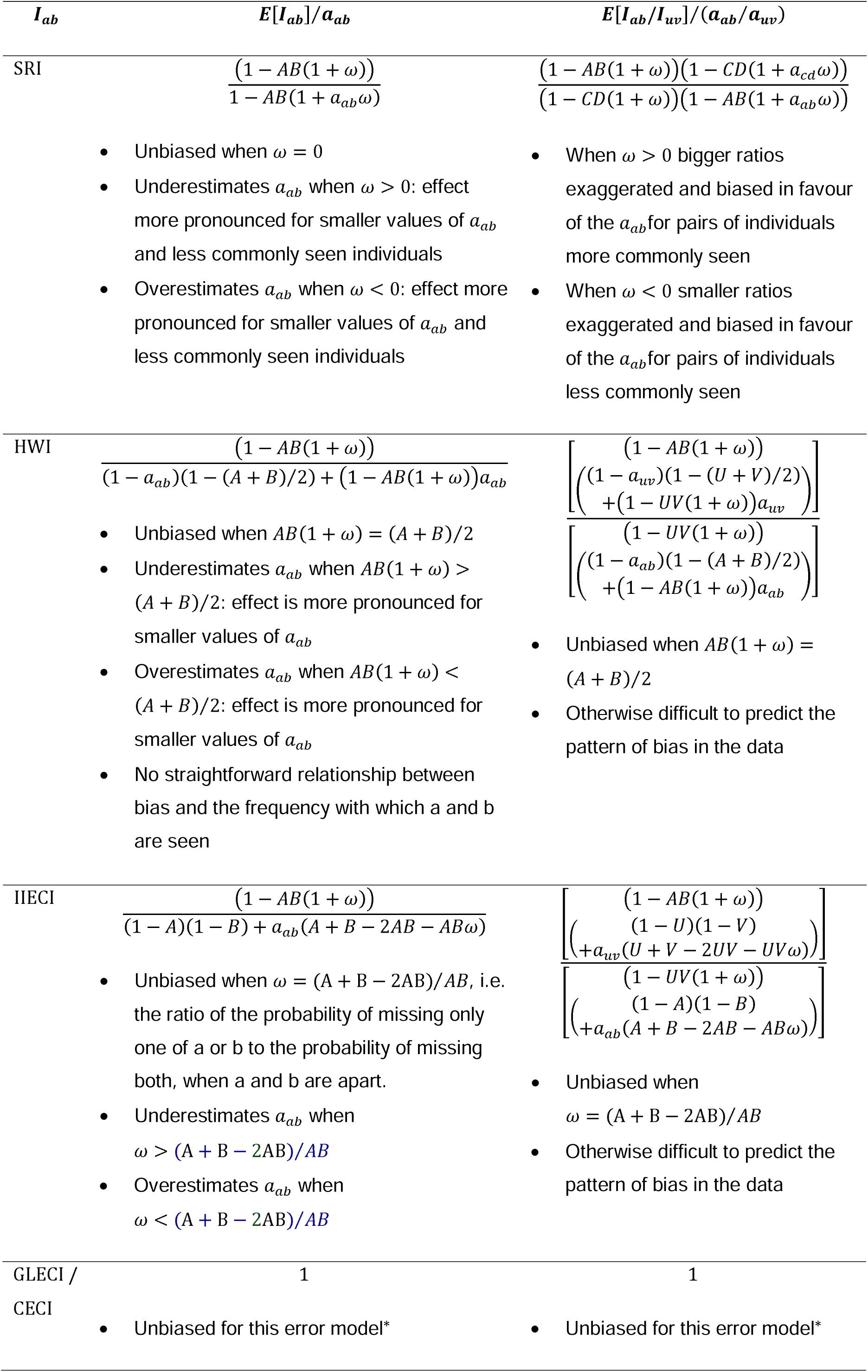
Biases in different association measures arising from group location error. *signifies no bias inferred from simulations.

The vSRI performs very poorly when only group location error is present, overestimating *a_ab_* whilst *ω* < (A + B – 2AB*)/AB.* This is because it excludes *y_a_* and *y_b_* from the denominator on the assumption that these data are uninformative, whereas in this scenario these are cases where we know that a and b were not associating. As *ω* increases, an increasing number of cases where a and b were associating are erroneously assigned to *y_null_.* Thus, exclusion of *y_null_* from the index eventually offsets the positive bias (when *ω* = (A + B — 2AB*)/AB)* resulting from exclusion of *y_a_* and *y_b_* from the denominator. The vSRI is therefore not a useful index as it contains assumptions that are unlikely to be met in the majority of studies.

The GLECI is generally unbiased because *y_null_* is included in the index in such a way that excludes this positive bias. Importantly, the 95% confidence intervals for the GLECI contained the true value of *a*_ab_ in close to 95% of cases, showing they perform validly in this scenario (see Fig 1b). In contrast, the 95% confidence intervals associated with the simple ratio, half weight index and IIEC index only performed acceptably for a very narrow range of values of *ω*. Furthermore, the GLECI (and CEI index with *ϕ=* 1) is the only index of those considered that is unbiased when estimating the ratio of two associations (see Table 1). This suggests that the SRI, HWI and IIECI are not suitable for estimating either the relative or absolute strength of associations when group location error is believed to be present, and that, ideally the GLECI should be used if calibration data can be obtained.

Note that the simulations assumed that the researcher has an accurate estimate of *ω* with which to calculate the GLECI (see below). In reality the better the estimate of *ω* is, the better the estimate of *a_ab_* will be, but even a rough estimate of *ω* will be preferable to no calibration at all. Furthermore, we assume that the probability of missing *a* and *b* when they are together will be (1 + *ω*) x that of missing both *a* and *b* when they are separate (*C* = (1 + *ω*)*ΑΒ*). Further work may conclude that this relationship does not generally hold, in which case the GLECI might be suitably modified to use a different calibration statistic. Nonetheless, the relationship posited here requires weaker *a priori* assumptions to be made about the data than the commonly used half weight index, which assumes that *C* = (A + B)/2. We show that when this assumption is even slightly wrong, the half weight index will be a poor estimate of *a_ab_.*

One possible option to resolve the half weight index is to generalize it to be an M weight index (MWI): *x/*(*M*(*y_a_* + *y_b_*) + *y_ab_ + x*). The M weight index assumes that the probability of seeing a and b together is equal to M x of the sum of the probability of seeing each of them each apart. Equivalently, missing *a* and *b* when they are together is M x more likely than missing them both when they are apart. Thus the MWI could be calibrated to the data analogously to the GLECI, but each index assumes a different relationship among the observation errors in the population.

When calibration data cannot be obtained, there is a strong case for preferring the simple ratio index. Use of the SRI results in biases that are more likely to be qualitatively predictable in the absence of calibration data, than biases resulting from the HWI or IIECI (see Table 1). For instance, when a researcher suspects that, in general, missing two individuals when they are together is more likely than missing both when they are apart, they can expect i) SRI values to be underestimates of *a_ab_;* ii) for this underestimation to be more pronounced for smaller values of *a_ab_* and for less commonly seen individuals; and iii) for bigger ratios between real associations to be overestimated relative to smaller ratios. Thus a researcher can assess whether these inaccuracies are likely have any great bearing on their conclusions in their specific case. In contrast, using the HWI we cannot easily make such qualitative predictions unless we are in a position to judge the relative size of *AB*(1 + *ω*) versus (*A + B*)/2; and, for the IIECI, *ω* versus (A + B – 2AB)/*AB*. Making such judgments empirically is likely to be at least as challenging as acquiring the calibration data required calculating the GLECI or a calibrated MWI. Consequently, if only group location error is present and calibration data cannot be obtained, we recommend use of the SRI, with careful consideration of how the biases identified above might affect the interpretation of the study.

### Individual identification error only

When only individual identification error is present we find that the IIECI is an unbiased estimator of *a_ab_*, whereas the simple ratio and half weight indices are biased downwards whenever ∈ *>* 0 (see Table 2 and Figure 2). This is because as *e* increases, more and more cases where a and b are associating are erroneously attributed to *y_a_* or *y_b_.* Since *y_a_* and *y_b_* are included in the denominator for the SRI and HWI, this results in an under-estimation of *a_ab_.* The effect is reduced in the HWI since *y_a_* and *y_b_* have a reduced weighting in this index. However, this is not sufficient to ensure that the HWI is valid for even small individual identification error rates. The GLECI was not included separately in these simulations since in this scenario (1 + *ω*) ∈^2^ = ∈^2^, so *ω* = 0, meaning the GLECI reduces to the simple ratio. Consequently, we can see that the GLECI also performs badly at estimating absolute association values when there is individual identification error but no group location error.

**Table 2:** Biases in different association measures arising from individual identification error

| $I_{ab}$ | $E[I_{ab}]/a_{ab}$ | $E[I_{ab}/I_{uv}]/(a_{ab}/a_{uv})$ |
| --- | --- | --- |
| SRI | $1 + \epsilon(2 - \epsilon)$ | 1 |
| HWI | $1 + \epsilon$ | 1 |
| IIECI/CECI | 1 | 1 |
| GLECI | $1 + \epsilon(2 - \epsilon)$ | 1 |

**Figure 2:**
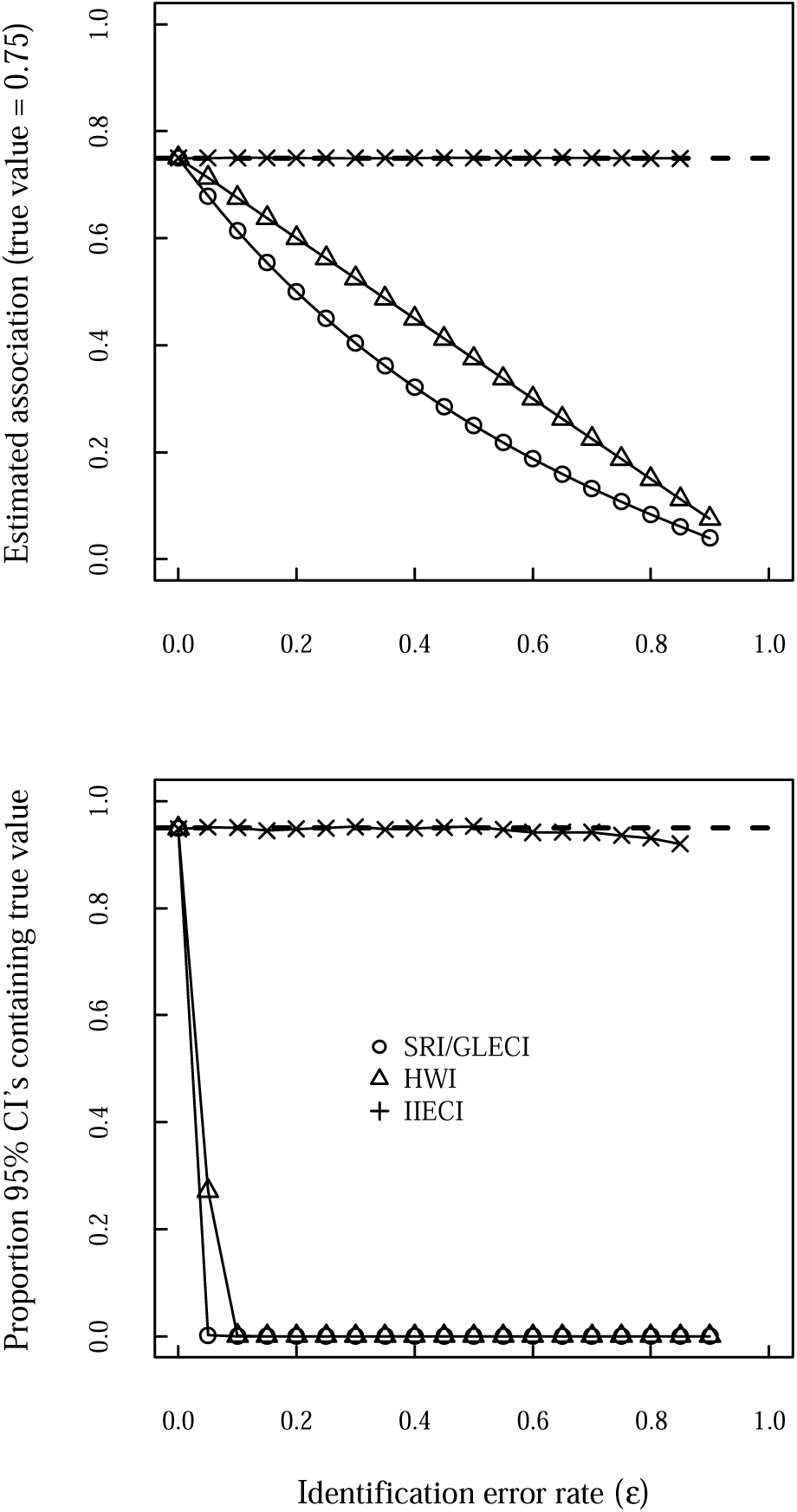
a) Bias in different association indices as a function of individual identification error (*ω*) when applied to simulated data; b) performance of 95% Wald confidence intervals as a function of *ω*. Similar results were obtained for a true association value of 0.25 and 0.75.

The 95% confidence intervals for the vSRI contained the true value of *a_ab_* in close to 95% of cases, showing they perform validly under a scenario of only individual identification error (see Fig 2b). This dropped slightly with very high error rates, as a result of the effective sample size decreasing as most data is attributed to *y_a_, y_b_* or *y_null_* (not that with small sample sizes Wald confidence intervals are likely to be anti-conservative: too narrow). In contrast, the 95% confidence intervals associated with the SRI (and hence the GLECI) and HWI performed very badly with even small individual identification error rates. Consequently, if a researcher is interested in estimating the absolute values of *a_ab_* and only individual identification error is likely to be present (we anticipate this scenario to be rare), we recommend use of the vSRI, which requires no calibration data.

In contrast to the group location error only, all association indices gave unbiased estimates of the relative size of associations between pairs of individuals (see Table 2). This means that if the research aims are purely in the scale free properties of a system, such as the relative position of individuals in a social network (Aplin, Firth, et al., 2015; Wilson, Krause, Dingemanse, & Krause, 2013), any of the indices considered will be sufficient. However, if there is also a risk of group location error, and appropriate calibration data cannot be obtained (see next section), we recommend use of the SRI to estimate relative associations due to the advantages in interpreting this index in the presence of group location error (see above).

### Combined errors

We find that the CECI is an unbiased estimator of *a_ab_* across the possible range of values for ∈, *ω* and *ϕ* when we assumed ∈_*a*|!*ab*_=∈ for all individuals (see Fig S1 in ESM). Furthermore, the CECI was also an unbiased estimator of *a_ab_* when ∈_*a*|!*ab*_ was allowed to vary across the population regardless of the magnitude of variation in ∈_*a*|!*ab*_ (see Fig S2 in ESM). In contrast, the SRI, HWI, IIECI and GLECI were biased in a manner that was dependent on the combination of values for ∈, *ω* and *ϕ.* When *ϕ* was close to 1, the pattern of bias was similar to when only group location error was present. In other words, when it is unlikely that only one of *a* and *b* will be missed when they are in the same group, bias is similar to when we have only group location error. As *ϕ* became smaller (more likely that only one of *a* and *b* will be missed when they are in the same group) all four indices start to underestimate *a_ab_* at a lower value of *ω*. The effects of both *ϕ* and *ω* are magnified more as ∈ gets larger. Consequently, if individual identification errors are likely to be common in addition to group location error, we suggest calibration data is acquired to estimate ∈, *ω* and *ϕ* and the CECI is used. The pattern of bias in the other indices will be difficult to predict qualitatively unless the risk of individual identification error is known to be small. Consequently, if calibration data cannot be obtained under such circumstances we suggest extra efforts are made to minimise individual identification error, and the SRI be used with the understanding that it will provide noisy estimates of *a_ab_.*

When we assume observation error is homogeneous across the population, the 95% confidence intervals for the CECI tend to contain the true value of *a_ab_* in >95% of cases (see Fig S3). This suggests that, at large sample sizes at least, the Wald confidence intervals are slightly too wide. However, given that Wald confidence intervals are always an approximation and are widely used in statistics, this is a minor concern. However, when observation error varied greatly across the population, the 95% confidence intervals became far too narrow (see Fig S4), as a result of the extra uncertainty that is unaccounted for in the derivation of the standard error. Correcting the standard errors for this uncertainty does not seem straightforward, though further work could address this if the CECI proves to be useful and becomes widely adopted. Therefore, we suggest that the standard errors and confidence intervals for the CECI be trusted as approximately valid if the variation in observation rate is believed to be small, and not be trusted if that variation is believed to be large.

Our recommendations for the choice of association index are shown as a flowchart in Fig. 3. The indices and their standard errors are shown in Table 3.

**Figure 3:**
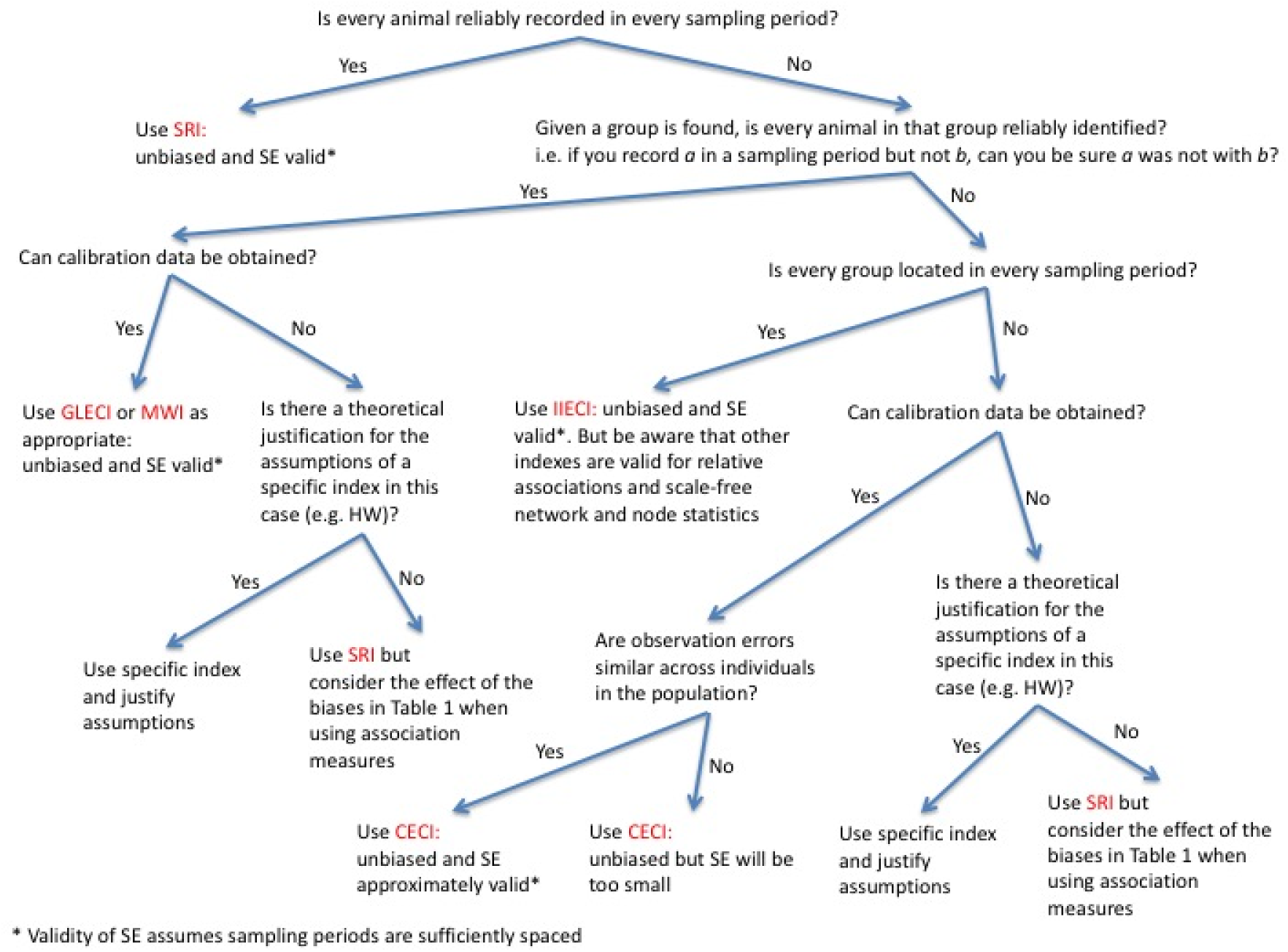
Flowchart with our suggested strategy for selecting an association index.

**Table 3:**
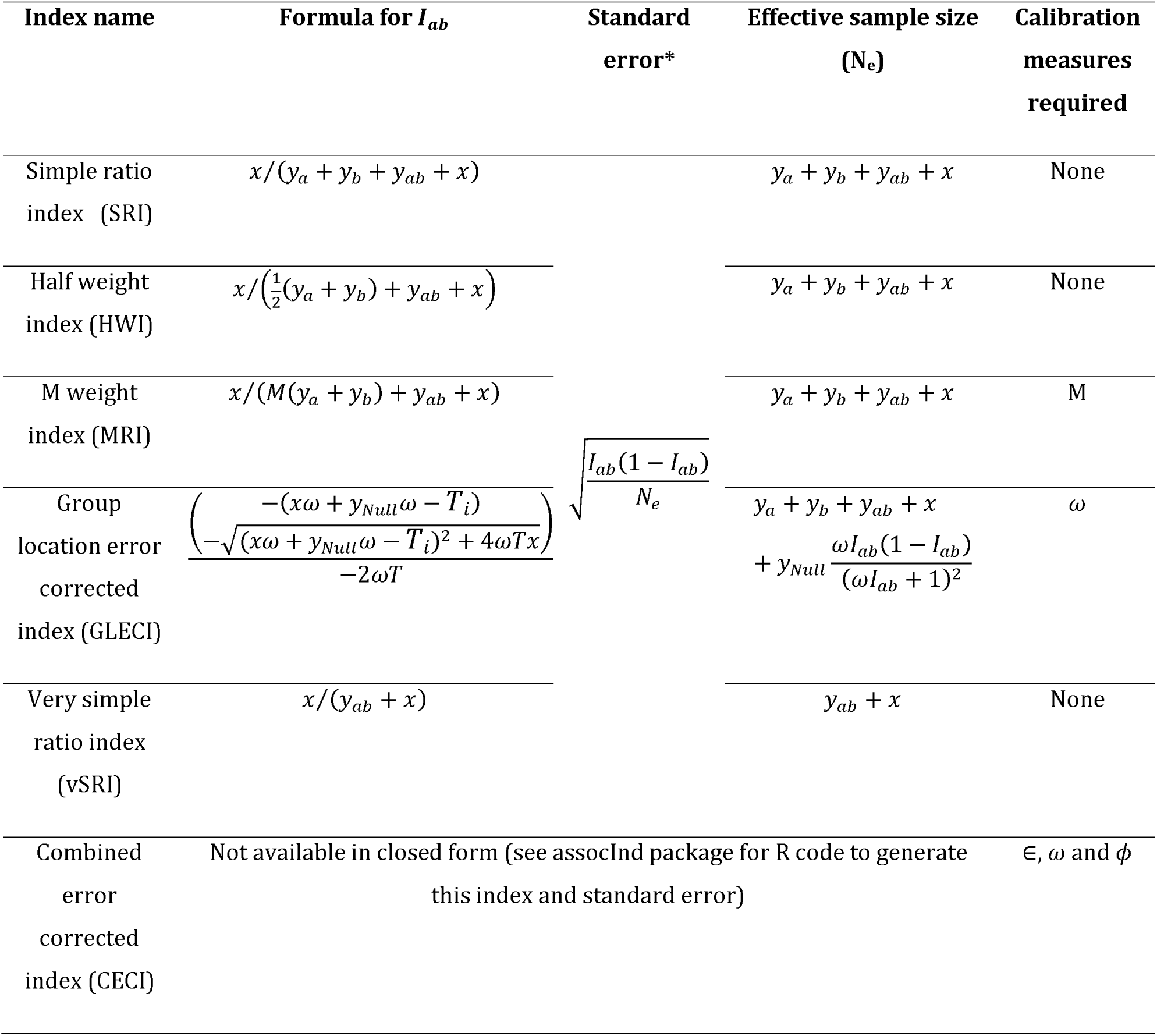
Summary of the indices considered in this paper. *Standard errors presented assume sampling periods are sufficiently spaced to be considered approximately independent, and that any calibration measures are known without error. See Supplementary Information for derivations of standard errors.

## OBTAINING CALIBRATION DATA

Here we suggest some initial ideas for obtaining calibration data that can be used to estimate the calibration parameters derived above. These suggestions can almost certainly be improved upon, by taking account of what data collection protocols are feasible in specific circumstances perhaps by deriving maximum likelihood estimates of calibration parameters given the data yielded by each such protocol. Here we limit ourselves to providing relatively simple intuitive ways of estimating calibration parameters. Whilst not optimal, our analysis above suggests that these methods are nonetheless likely to be an improvement on the unsupported use of a non-calibrated index such as the HWI.

One way a researcher might obtain estimates of calibration parameters is to collect data that can be assumed to be approximately error free for a subset of individuals, whilst simultaneously collecting association data using their standard protocol. This could be done by focal follows of a sample of individuals conducted by one researcher, whilst another collects data using the association data collection protocol. Alternatively some individuals could be tagged with GPS or proximity loggers (Kays, Crofoot, Jetz, & Wikelski, 2015; Krause et al., 2013) able to record encounters between individuals with more precision.

First we suggest that a researcher assess whether or not individual identification error is present and important. This could be done by calculating the proportion of sampling periods in which each individual included in the error free dataset was present in a group that was located using the standard protocol but not recorded as being present. If this proportion is 0 or close to 0 for most individuals, we suggest individual identification error be ignored, and the GLECI or MWI can be used. Otherwise, the CECI should be used (unless group location error is believed not to be present, in which case the vSRI should be used, which does not require calibration).

If individual identification error is not present or negligible, the researcher needs to choose between the GLECI and MWI and then estimate the relevant calibration parameter (*ω* or m). For any two individuals *a* and *b* in the error free sample, we know in which sampling periods they were together and in which they were not together. This enables the researcher to calculate *y_null|ab_*, the number of times *a* and *b* were not recorded by the association protocol during the calibration data collection period, when a and b were known to be together. *C*, the probability *a* and *b* will be missed when they are together, can be estimated as *y_null|ab_/N_ab_* where *N_ab_* is the number of sampling periods that *a* and *b* were known to be together. A can then be estimated as the proportion of sampling periods in which *a* was not recorded by the association protocol and known not to be with *b.* B can be estimated in an analogous manner.

This process can be repeated for every combination of two individuals in the error free sample. The researcher can then use plots of these data to choose between the GLECI and MWI. If the assumptions of the GLECI hold, we would expect C to have a linear relationship with AB, with a slope of (1 + *ω*), whereas if the assumptions of the MWI hold, we would expect C to have a linear relationship with (A + B) with a slope of 1/m. We suggest the researcher make each plot to decide which assumption is most realistic, and thus chose between the GLECI and MWI. If the GLECI is chosen, they can then fit a linear regression (constrained to pass through the origin) of C against AB and take *ω* = slope-1. If the MWI is chosen, they can then fit a linear regression (constrained to pass through the origin) of C against A + B and take m = 1/slope.

The CECI requires estimates of *ω, ϕ* and ∈. ∈ can be estimated as the population average of ∈_*a*|!*ab*_. To this end, we suggest for each dyad of individuals *a* and *b*, where *a* is an individual for which we have error free data, we calculate *M_a|_*_!*ab*_ the number of sampling periods in which the standard sampling protocol missed individual *a* and we know (from the error free sample) that *a* and *b* were not together. We can then estimate ∈_*a*|!*ab*_ = *M_a|_*_!*ab*_/*N_!ab_*, where *N_!ab_* is the number of sampling periods that *a* and *b* were known not to be together. ∈ can then be estimated as the average of both ∈_*a*|!*ab*_ across all dyads containing at least one individual in the error free dataset. Next the researcher can estimate *ω*, using the relationship ∈_*ab*|*ab*_ *=* (1 + *ω*) ∈_*a*|!*ab*_ ∈_*b*|!*ab*_ in the same manner as suggested for estimating *ω* for the GLECI above.

To estimate *ϕ* a researcher can use the relationship:

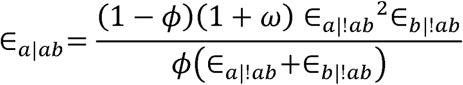

For each dyad of individuals for which we have error-free data a researcher can obtain an estimate of ∈_*ab*|*ab*_ =*M_a|ab_/N_ab,_* where *M_a|ab_* is the number of sampling periods in which the standard sampling protocol missed individual *a* and we know (from the error free sample) that *a* and *b* were together, and where *N_ab_* is the number of sampling periods that *a* and *b* were known to be together. Using the estimates of ∈_*a*|!*ab*_ and ∈_*b*|!*ab*_ obtained above, the researcher can then obtain an estimate of ∈_*a*|!*ab*_ ^2^∈_*b*|!*ab*_/ (∈_*a*|!*ab*_ + ∈_*b*|!*ab*_). ∈_*b*|!*ab*_ can be likewise be estimated as *M_b|ab_/N_ab_.* and ∈_*b*|!*ab*_^2^ ∈_*a*|!*ab*_^2^/ (∈_*a*|!*ab*_ + ∈_*b*|!*ab*_) estimated as for individual *a.* The researcher can then fit a linear regression (constrained to go through the origin) with the set of ∈_*a*|*ab*_ and ∈_*b*|*ab*_ as the dependent variable and the set of ∈_*a|*!*ab*_^2^ ∈_*b*|!*ab*_ /(∈_*a|*!*ab*_ + ∈_*b|*!*ab*_) and ∈_*b|*!*ab*_^2^∈_*a|*!*ab*_ /(∈_*a|*!*ab*_ + ∈_*b|*!*ab*_) as the independent variable. Since the slope of the regression will estimate (1 – *ϕ*)(1 + *ω)/ϕ*, we can estimate *ϕ* = (1 + *ω*)(*slope* + 1 + *ω*).

A second way we suggest a researcher might obtain estimates of calibration parameters is to have two researchers or research teams independently collecting association data using their standard protocol, for a portion of the data collection period. For example, this might be done by having the second researcher (denoted *Y)* collect data a short time after the first (denoted *X*), on a short enough time scale that group composition is unlikely to have changed. Here we suggest a procedure for obtaining calibration statistics for the CECI since this reduces down to the IIECI or GLECI when the calibration data reveals the relevant component of error to be absent.

We suggest that researchers first obtain estimates of ∈_*a|*!*ab*_. Ideally we wish to estimate *M_Xa_*_|!*ab*_/*N_!ab,_* the proportion of events that X missed *a* given *a* and *b* were together. We suggest researchers do this by calculating the proportion of sampling periods *X* missed *a* given that *Y* recorded *a* and *b* in different groups (which we denote *Y!ab*), i.e. *M_Xa|Y!ab_/N_Y!ab_.* We can repeat this procedure, reversing the role of *X* and *Y* to obtain *M_Ya|X!ab_/N_X!ab_.* We can then take the mean as our estimate, i.e. ∈_*a|*!*ab*_ = *M_Xa|Y!ab_/2N_Y!ab_* + *M_Ya|X!ab_ /2N_X!ab_.* ∈ can then be estimated as the population average of ∈_*a|*!*ab*_ as above.

Researchers can then obtain estimates of ∈_*a|ab*,_ the probability that individual *a* is missed when it is in a group with individual *b.* Again, we have potentially two estimates of this for each combination of *a* and *b.* First we have *M_Xa|Yab_/N_Yab,_* the proportion of sampling periods in which researcher X missed individual *a* given researcher Y recorded *a* and *b* together, and conversely we have *M_Ya|Xab_/N_Xab_.* We suggest ∈_*a|ab*_ be estimated as the average of these, i.e. ∈_*a|ab*_ = *M_Xa|Yab_/2N_Yab_* + *M_Ya|xab_/2N_Xab_.* We suggest this be done for all combinations of *a* and *b* for which *a* and *b* were frequently seen together, in order to obtain estimates of *ω* and *ϕ. ω* can be estimated using the relationship ∈_*a|ab*_ = (1 + *ω*) ∈_*a|!ab*_ ∈_*b|!ab*_ in the same manner as suggested for estimating *ω* for the GLECI above, *ϕ* can be estimated as above by fitting a linear regression (constrained to go through the origin) with the set of ∈_*a|ab*_ and ∈_*b|ab*_ as the dependent variable and the set of ∈_*a|!ab*_ ^2^∈_*b|!ab*_/(∈_*a|!ab*_ + ∈_*b|!ab*_) and ∈_*b|!ab*_ ^2^ ∈_*a|!ab*_ /(∈_*a|!ab*_ + ∈_*b|!ab*_) as the independent variable yielding the estimate *ϕ* = (1 + *ω*)/(*slope* + 1 + *ω*).

## DISCUSSION

Studies of animal social networks have shed new light on many ecological and evolutionary processes. For example, the structure of the social environment can shape how information (Aplin, Farine, et al., 2015; Aplin, Farine, Morand-Ferron, & Sheldon, 2012; Farine, Aplin, Sheldon, & Hoppitt, 2015) and diseases (K.L. VanderWaal, Atwill, Isbell, & McCowan, 2013; K. L. VanderWaal et al., in press) spread in wild populations. Further, they have provided important insights into the role of the social environment on shaping selection (Farine & Sheldon, 2015; Formica et al., 2011; McDonald, 2007; Oh & Badyaev, 2010; J.B. Silk, Alberts, & Altmann, 2003; J. B. Silk et al., 2010; Wey, Burger, Ebensperger, & Hayes, 2013). However, studies have used varying approaches to quantify the relationships among individuals. Whilst care is generally taken to ensure that the chosen approach has biological relevance, the underlying assumptions behind the approach used are almost never explicitly considered. The results of our study into different association indices suggests that many commonly-used approaches should be avoided as they do not accurately estimate the (absolute or relative) strengths of social bonds, which has implications on estimates of social structure and social processes occurring through social networks.

Ideally, studies of animal social networks would capture information about all individuals in the study population at once. Realistically, this is unlikely to be possible in all but a very select number of studies. Thus, before constructing a social network from a given set of data, we suggest that the following questions should be addressed:

1. How much data has been collected on each dyad?
2. Are all individuals sampled equally?
3. What proportion of the population is observed in each sample?
4. Are there any mistakes in the observations?

The issues surrounding question 1 have now been relatively well outlined in the literature (Farine &Strandburg-Peshkin, 2015; Franks et al., 2010; Lusseau, Whitehead, & Gero, 2008; M. J. Silk et al., 2015; Whitehead, 2008). In general, these studies have found that collecting enough data on each dyad (at least 20 observations per dyad) is important for accurately estimating global social network structure. In the current study, we address issues arising from questions 2 and 3, and how better indices can reduce the potential impact that missing observations can have on both the absolute and relative estimates of association strengths among individuals. Question 4 represents an area requiring some further investigation.

Our study suggests that a critical step in the study of animal social networks will be the collection of calibration data. Currently-used association indices are all based on arbitrary rates of missing observations. For example, the half-weight index assumes that the probability of missing individuals a and b when they are together is exactly half the probability of missing either individual when they are apart. Importantly, we have shown that when this is not true, the HWI does not result in a ‘better approximation’ of the real association rate when compared to the simple ratio index. Thus, we recommend avoiding the use of the HWI, and instead using the SRI when no calibration data is available (see Figure 3). In reality, it is likely that the rates of observation could be estimated from parameters of the observation data, such as the average group size, the average number of individuals observed in a sampling period, and the average number of groups sampled. Whether these can be used to parameterize the M-weighted index warrants further investigation.

Several extensions of association indices have been proposed to deal with other issues arising when sampling populations. Godde, Humbert, Cote, Reale, and Whitehead (2013) suggest a method to correct for the fact that individuals that prefer large groups are more likely to be observed together. This involves normalizing the association index values by the two individual’s combined gregariousness (the sum of their association indices to others). To deal with other potentially confounding influences on association patterns (such as home-range overlap) when attempting to estimate true association rates, Whitehead and James (2015) propose regressing association indices against other input parameters. Our proposed indices work equally well with both of these approaches as they are simply new ways of defining the association value for pairs of individuals. The above two studies highlight how patterns of affiliation can be affected by a range of different factors. Thus, even if good calibration data can be obtained to estimate accurate relationship strengths, it will always be important to use null models when conduction hypothesis testing with animal social networks (Farine *in review*).

We encourage further investigation into methods for collecting informative calibration data alongside the social network data. One potential avenue could be to use mark-recapture techniques that explicitly investigate detection probabilities, and these could be conditioned on having observed one or more particular individuals. There are also increasing numbers of studies that are collecting complete datasets from groups or populations of animals, and these could provide very useful data for testing different approaches to collect calibration data. A particular challenge that will arise is that social network analysis has proved particularly useful in species or communities that exhibit fission-fusion dynamics (Aureli et al., 2008; Couzin, 2006; M. J. Silk, Croft, Tregenza, & Bearhop, 2014). Here, the rate of turnover in group membership can be very rapid (for example group membership in great tits, Parus major, can be close to random after just 10 minutes,Farine, Firth, et al., 2015). Similarly, when using focal observations—following a single individuals and recording its interactions with others—the concept of a ‘group’ is unclear, and we do not yet have a definition for out to estimate the group location error. This is because all individuals can be observed, but only interactions among a subset of edges (those connected to the focal) are recorded. Thus, in such systems, more research is required to find robust approaches for collecting calibration data.

In all of our simulations, and more generally in the assumptions of how well any analysis captures reality, our estimates of accuracy are a ‘best-case scenario’. In reality, most datasets will also contain erroneous observations. At best, these are simply individuals wrongly assigned into a group in which they did not occur. If such individuals (and their erroneous associates) are observed many times, then the resulting association strength will be low, and the error will be reasonably well dealt with by using weighted social networks. However, the impact of incorrectly assigning one individual identity for another could be significantly greater, especially as such errors are unlikely to be randomly distributed throughout the dataset (i.e. certain pairs of individuals are more likely to be confounded than others). No association index will be able to correct for such errors. However, the relative effect that incorrect assignments of identity have on different approaches to estimate social network structure warrants further investigation.

Our paper makes it clear that we should avoid blindly using association indices without proper consideration of the assumptions that they entail. We also recommend discontinuing the use of the HWI, which Cairns and Schwager (1987) already 30 years ago identified as being problematic because of the many assumptions it makes. Instead, we recommend using properly calibrated association indices wherever possible, and using the SRI if no appropriate calibration data or estimates of rates of detectability are available. Whichever approach is used, we hope that our paper will at least encourage researchers to carefully and explicitly consider their choice of approach for estimating association strengths among individuals in their study population.

## ACKNOWLEDGEMENTS

DRF received additional support from the BBSRC (BB/L006081/1 to B.C. Sheldon).

